# Embodiment of Supernumerary Robotic Limbs in Virtual Reality

**DOI:** 10.1101/2022.01.31.478373

**Authors:** Ken Arai, Hiroto Saito, Masaaki Fukuoka, Sachiyo Ueda, Maki Sugimoto, Michiteru Kitazaki, Masahiko Inami

## Abstract

The supernumerary robotic limb system expands the motor function of human users by adding extra artificially designed limbs. It is important for us to embody the system as if it is a part of one’s own body and to maintain cognitive transparency in which the cognitive load is suppressed. Embodiment studies have been conducted with an expansion of bodily functions through a “substitution” and “extension.” However, there have been few studies on the “addition” of supernumerary body parts. In this study, we developed a supernumerary robotic limb system that operates in a virtual environment, and then evaluated whether the extra limb can be regarded as a part of one’s own body using a questionnaire and whether the perception of peripersonal space changes with a visuotactile crossmodal congruency task. We found that the participants can embody the extra-limbs after using the supernumerary robotic limb system. We also found a positive correlation between the perceptual change in the crossmodal congruency task and the subjective feeling that the number of one’s arms had increased (supernumerary limb sensation). These results suggest that the addition of an extra body part may cause the participants to feel that they had acquired a new body part that differs from their original body part through a functional expansion.

## Introduction

Research on supernumerary robotic limbs (SRL)^1^ aims to add new limbs as an extension of a body function. Several methods have been reported to manipulate a system^2^ by linking the movement of a certain innate limb (called limb mapping)^3,4^, linking to facial expressions^5^, or linking to myoelectric signals, i.e., an electromyogram (EMG)^6^. One of the prerequisites of the SRL system is the cooperative behavior between humans and systems, and the system is expected to be able to move voluntarily according to the intentions of the operator, just like with an innate limb.

It is important to be able to treat an SRL as if it is a part of one’s own body when constructing it as a new body part. In “*We feel well as long as we do. We feel well as long as we do not feel our body*”^7^, it is stated that if we do not pay attention to our bodies, we will not feel resistance to our body states and movements. In other words, if we can generate a state in which the cognitive workload during a body movement is suppressed (i.e., cognitive transparency is ensured), it can be interpreted that foreign parts can be treated as if they are our own bodies. In fact, when the cognitive workload is high, the quality and accuracy of the manipulation are hindered^8^. Considering the design of the SRL system, if the cognitive workload can be controlled, the system will work seamlessly for the operator and the system can be treated as part of the body. To the best of our knowledge, in conventional research on extra-limb robotics, whether the system can be treated as a part of the body has not been sufficiently investigated. This study is aimed at exploring this point.

In cognitive science, it is known that human perception can be transformed when it is sufficiently applied to the use of tools, and that tools can be treated as if they are a part of our body^9^. This is called “tool embodiment”^10,11^, which has been examined from the perspective of neuroscience and cognitive science^12–15^. Merleau-Ponty explained that the repeated use of a walking stick by a blind person not only incorporates the stick into the blind person’s body image, it also makes the stick a physical aid, which is an extension of bodily synthesis^16^. Body cognition against the embodiment of tools has been interpreted in various fields. When we consider it again from the perspective of cognitive science, it is important to know whether external tools can be incorporated into our body schema and body image. Body schema is a perceptual model generated from sensory signals, such as movement and posture, and is used to guide body movements and motions^17^. By contrast, body image is an internal model of the body constructed from visual information and is used to make perceptual judgments^18^. These models attempt to interpret body perception from both motor and perceptual perspectives and can be modified not only by the innate body but also by extrinsic tools and environmental influences^7,19^.

In the discussion of embodiment, the main explanatory variables and elements in the field of cognitive science are the sense of body ownership (SoO), sense of agency (SoA), and sense of self-location (SoSL). Gallagher defined minimal self as the smallest unit of self-consciousness and argued that it is composed of SoO and SoA^20^. A sense of body ownership refers to a state in which a person perceives and reacts emotionally to an object as if it is his or her own body^21^. This is one of the basic elements of the embodiment of objects and tools. The rubber hand illusion (RHI) is a typical research example of sense of body ownership^21–25^. This refers to the phenomenon in which the feeling of one’s own hand, which is visually hidden, is gradually substituted or transferred to a rubber hand placed beside it when the rubber hand is repeatedly touched with a brush, resulting in the individual feeling as if the rubber hand is his/her. By contrast, a sense of agency is a state in which one can feel that the result of an action is their own^26–28^. Furthermore, a sense of self-location has been proposed as an embodiment for avatars placed in immersive spaces, such as virtual reality (VR)^29^. Other types of perception have also been considered as components to explain an embodiment.

Peripersonal space (PPS) has also been examined as an explanatory variable or element of embodiment^30^. Peripersonal space refers to the space surrounding the vicinity of the body, where stimuli from the outside world can be directly perceived. Humans perceive it through the integration of multiple sensory modalities, such as vision, touch, and auditory stimuli. In addition, spatial representation in the brain is said to be three-dimensional and is known as higher-order cognitive perception^31–34^. Peripersonal space is also thought to occur within the vicinity of embodied tools^12,35,36^, and the relationship between peripersonal space and sense of body ownership against an object has been investigated^37^. In addition, peripersonal space and body schema are closely related^38^, and are thought to affect the transformation of body movements.

Substitution or extension perceptual changes have been reported in many studies in the context of tool embodiment, while several prior studies, though not as numerous, have reported perceptual changes of addition. For example, the supernumerary hand illusion, which is an extension of the rubber hand illusion, attempts to add an extra limb by adding a rubber hand or virtual arm and fingers^39–41^. In this experimental paradigm, the subject is presented with both his/her own hands and a rubber hand, and visual and tactile stimuli similar to those of the rubber hand illusion are given simultaneously to duplicate the perception. As a result, the subject can feel a sense of body ownership of the rubber hand without losing the sense of body ownership against his/her innate hand. In addition, neuroscience has confirmed cases in which patients perceive an extra limb even though no such limb exists, and central nervous system disorders have been reported to cause supernumerary phantom limbs^42^. As an example of handling additional body part capable of voluntary movement, there are previous studies that evaluate changes in body perception and human flexibility by giving a third arm^43,44^, a sixth finger^45,46^ or a tail^47^. To the best of our knowledge, in conventional research on the robotics of extra body part, whether the system can be treated as a part of the body has not been sufficiently investigated. This study is aimed at exploring this point, following earlier work by Sasaki et al.^3^ and Drogemuller et al.^44^. In the field of SRL, which aims to expand different functions by adding body parts, it has not been sufficiently evaluated whether humans can acquire body representation or peripersonal space, including supernumerary limbs.

In this study, we developed a SRL system that operates in a VR environment, and evaluated whether artificial supernumerary limbs can be regarded as a part of one’s own body and what kind of perceptual changes occur when wearing the system (see Figure1). We measured the reaction times involved in the Crossmodal Congruency Task (CCT) before and after a learning task in which the participants learned to manipulate an SRL. The CCT is a measure that has been widely used to investigate the integration of vision and touch in the peripersonal space^31,48^. With this task, participants make discriminative judgments regarding the presentation position (up or down) of tactile stimuli while ignoring task-irrelevant visual distractor. We used the CCT to evaluate whether a strong integration of visual and tactile sensations, which is a typical feature of body perception, occurs within the vicinity of the body space for the extra limb, as a crossmodal congruency effect (CCE) score. In addition, we collected subjective evaluations related to the embodiment of SRL through a questionnaire.

**Figure 1.**
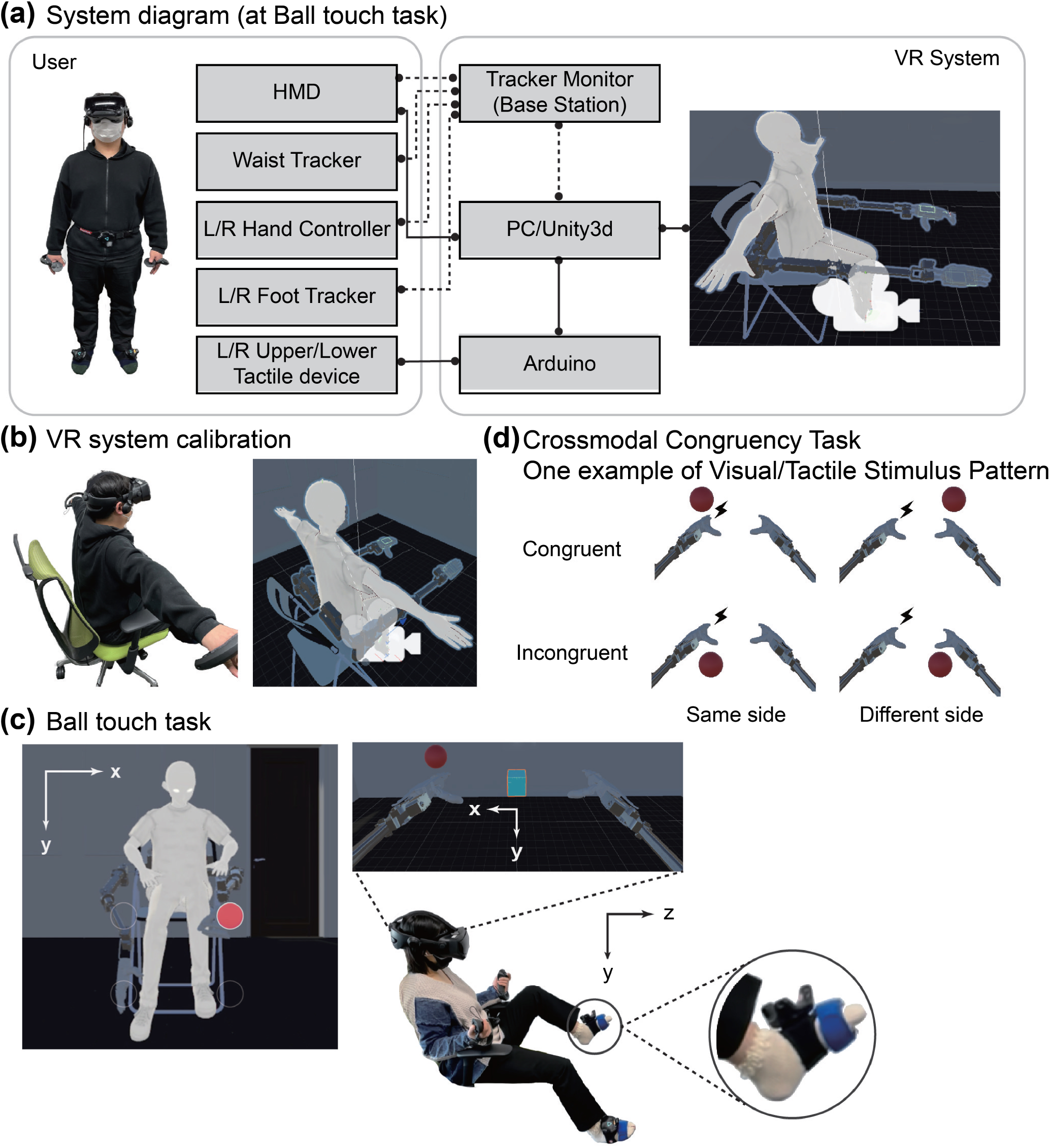
(a) The system diagram depicts the components of the system used in this experiment and their relationships. The solid line indicates a wired connection, and the dotted line indicates a wireless connection. (b) VR system calibration shows the correspondence between the real space and the VR space. (c) The first-and third-person perspectives of a ball touching task are shown. When the participant touches the ball, it vibrates to the location corresponding to the position of the innate foot. (d) An example of the visuotactile stimulus condition is a crossmodal congruency task (CCT). In this example, the tactile stimulus was transmitted to the back of the left foot, and the visual stimulus was presented to the back of the hand and palm of the left and right robot arms.

The results showed that the CCE scores were significantly different before and after the training, and a subjective evaluation showed that the participants had an embodied SRL. Surprisingly, there was a positive correlation between the changes in the CCE scores and the changes in the subjective evaluation scores of feeling that one’s arm had increased in comparison to the two innate arms before and after learning.

In this paper, we present the possibility of embodiment of the SRL system and the generation of peripersonal space, aiming at a functional expansion through the addition of body parts, although the evaluation was conducted in a VR environment; in addition, we report the possibility that the participants felt that they had acquired new body parts that were different from their own innate body parts. As a result, we suggest a direction of the design evaluation of the SRL system as well as the importance of discussing in detail the embodiment in cognitive science owing to the addition of the extra body part.

## Results

We focused on four measures for analysis in this experiment: (1) analysis of the learning task of the SRL system, (2) CCE score for reaction time collected in the CCT, (3) the questionnaire score of embodiment change for the SRL, and (4) the correlation between the CCE score and the questionnaire score of an embodiment change.

### Ball-touch task as learning of supernumerary robotic limbs manipulation

The subjects were asked to conduct a ball-touching task to adapt to the use of the SRL system in a VR environment. In this task, the subjects were required to touch a ball displayed randomly up, down, left, and right with the hand at the end of the supernumerary robotic arm, without setting any time limit. We prepared four sets of 100 touches per set, with a break at the end of each set. The average time required per set was analyzed to determine the tendency of the ball-touching task. To eliminate cases in which the ball appeared in the same place continuously and cases in which an excessive time was required for the task, cases in which it took less than 0.5 s or more than 10 s to complete the task were eliminated from the analysis. At the beginning of the task, it tended to take a long time to move the robot arm to touch the ball as intended, but it was confirmed that the participants became accustomed to the operation in the latter half of the task. Figure 2 shows a box plot of the duration time for each set of ball touches. As a result, the average time required to complete the task per set was 3.7 min (*±*0.8 min). The average time required for each set was 4.5 min (*±*1.1 min) for the first set, 3.6 min (*±*0.5 min) for the second set, 3.4 min (±0.4 min) for the third set, and 3.3 min (±0.3 min) for the final set. (0.4 min) for the third set, and 3.3 min (0.3 min) for the final set.

**Figure 2.**
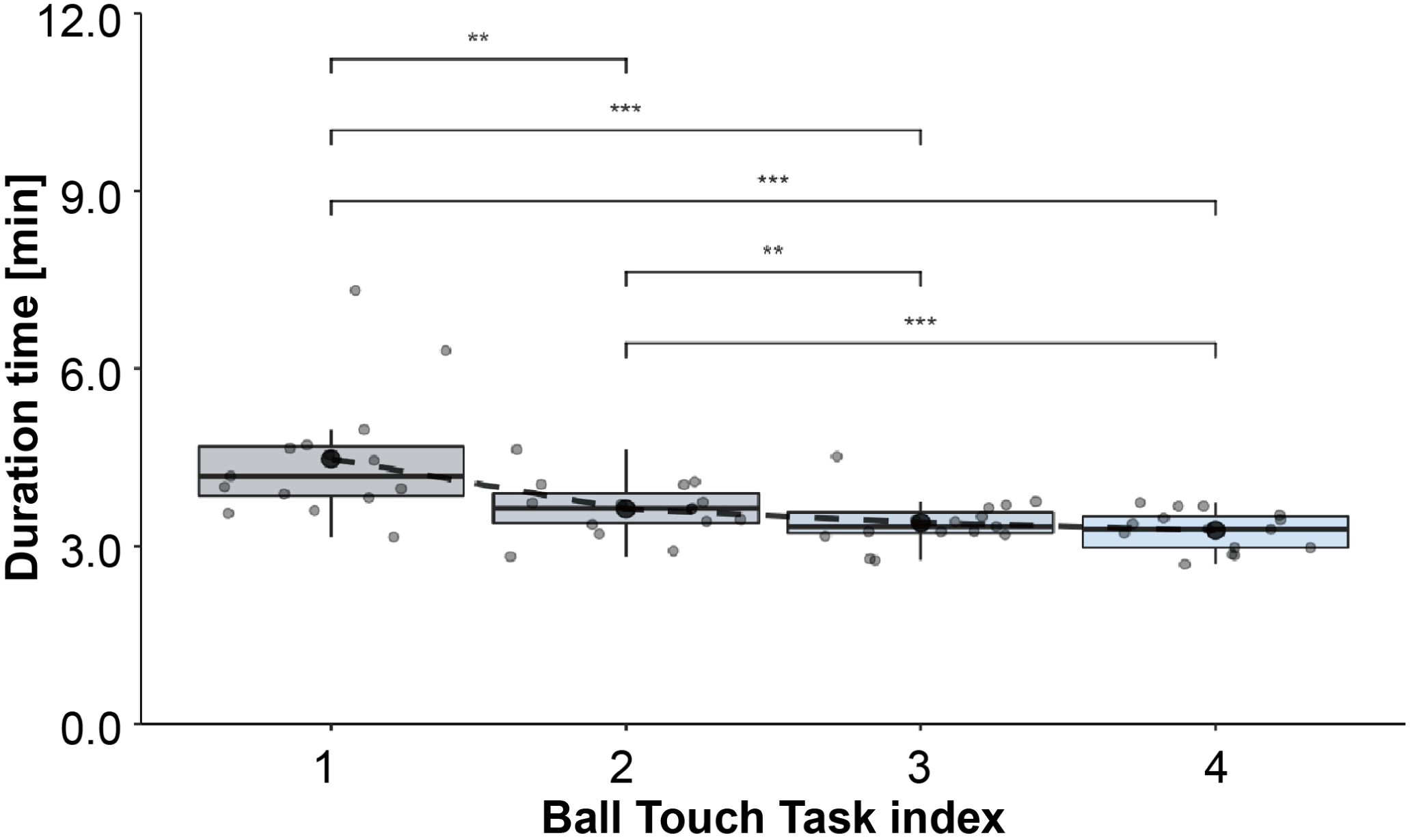
The duration time per set in the ball touch task. The circled markers are the results of the subjects, and the box plot shows the characteristics of the distribution of each set. * … *p <* 0.05, ** … *p <* 0.01, *** … *p <* 0.001

A Friedman test was applied to the mean time for each set, and a statistically significant difference [*χ*^2^(3) = 28.6, *p <* 0.001] was confirmed. In addition, the pairwise Wilcoxon rank-sum test using a Bonferroni correction showed statistically significant differences in all combinations except for comparisons between the third and fourth sets [adjusted p-value of “first set vs. second set” *<* 0.001, whereas those of “first set vs. third set” *<* 0.001, “first set vs. fourth set” *<* 0.001, “second set vs. third set” *<* 0.001, “second set vs. fourth set” *<* 0.001, and “third set vs. fourth set” = 0.81 *>* 0.05].

### Crossmodal Congruency Task

The CCT was used to evaluate whether a strong integration of visual and tactile sensations, a typical feature of body perception, occurred in the peripersonal space around the supernumerary robotic arms before and after the learning task. As task-irrelevant visual distractor was presented within the vicinity of the supernumerary robotic arms, the reaction time (RT) and accuracy of the responses to tactile stimuli presented to the toes were collected as data for analysis. To take into account the fact that some responses may be incorrect, the inverse effect (IE), which is the reaction time (RT) divided by the accuracy ratio of correct responses for each condition, was used for a statistical analysis and CCE score calculation^34,49^. Trials with reaction times of greater than 1500 ms were excluded as operational errors based on previous studies^48^.

The mean values of the inverse effect-based reaction time (IE-RT) were analyzed using a within-subject three-way repeated measure ANOVA, where the three factors of the ANOVA design were the vertical congruency of the stimulus presentation position (*Congruent* vs. *Incongruent*), the left-right lateral congruency of the stimulus presentation position (*Same* vs. *Different*), and the *pre-* and *post-*learning of the system. The results showed a main effect of vertical congruency (*Congruent* vs. *Incongruent*) [*F*(1, 14) = 46.797, *p <* 0.001, 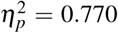] and an interaction of the three factors [*F*(1, 14) = 4.907, *p* = 0.044 *<* 0.05, 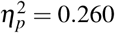].

Because a three-factor interaction was found in the mean IE-RTs, post-hoc analyses were conducted to examine the inverse effect-based crossmodal congruency effect (IE-CCE) scores and the effects observed between *pre-* and *post-*learning. The IE-CCE score can be calculated from the difference in IE-RT between the vertical *incongruent* and *congruent* trials. A two-way repeated measure ANOVA was applied to the mean of the IE-CCE scores, and the two factors in the ANOVA design were left-right lateral congruency of the stimulus presentation position (*same*/*different*) and *pre-*/*post-*learning required to wear the system. The results confirmed the main effect of the *pre-*/*post-*learning provided to wear the system [*F*(1, 14) = 6.237, *p* = 0.026 *<* 0.05, 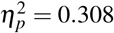] and the interaction of the two factors [*F*(1, 14) = 6.823, *p* = 0.021 *<* 0.05, 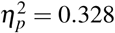] was confirmed.

Because a two-factor interaction was observed in the mean IE-CCE scores, we conducted multiple comparisons between six pairs consisting of four combinations (*pre-same, post-same, pre-diff* and *post-diff*) by adjusting p-value using the Bonferroni method^50^. The results (Figure 3) showed statistically significant differences in the IE-CCE scores of *pre-*/*post-*learning [*ad j*.*p* = 0.027 *<* 0.05] and of *same-*/*different-*side after learning [*ad j*.*p* = 0.013 *<* 0.05]. No statistically significant differences were found for the other conditions. This means that the IE-CCE scores increased significantly after learning to use the supernumerary robotic limbs only when the tactile stimulus was presented ipsilaterally.

**Figure 3.**
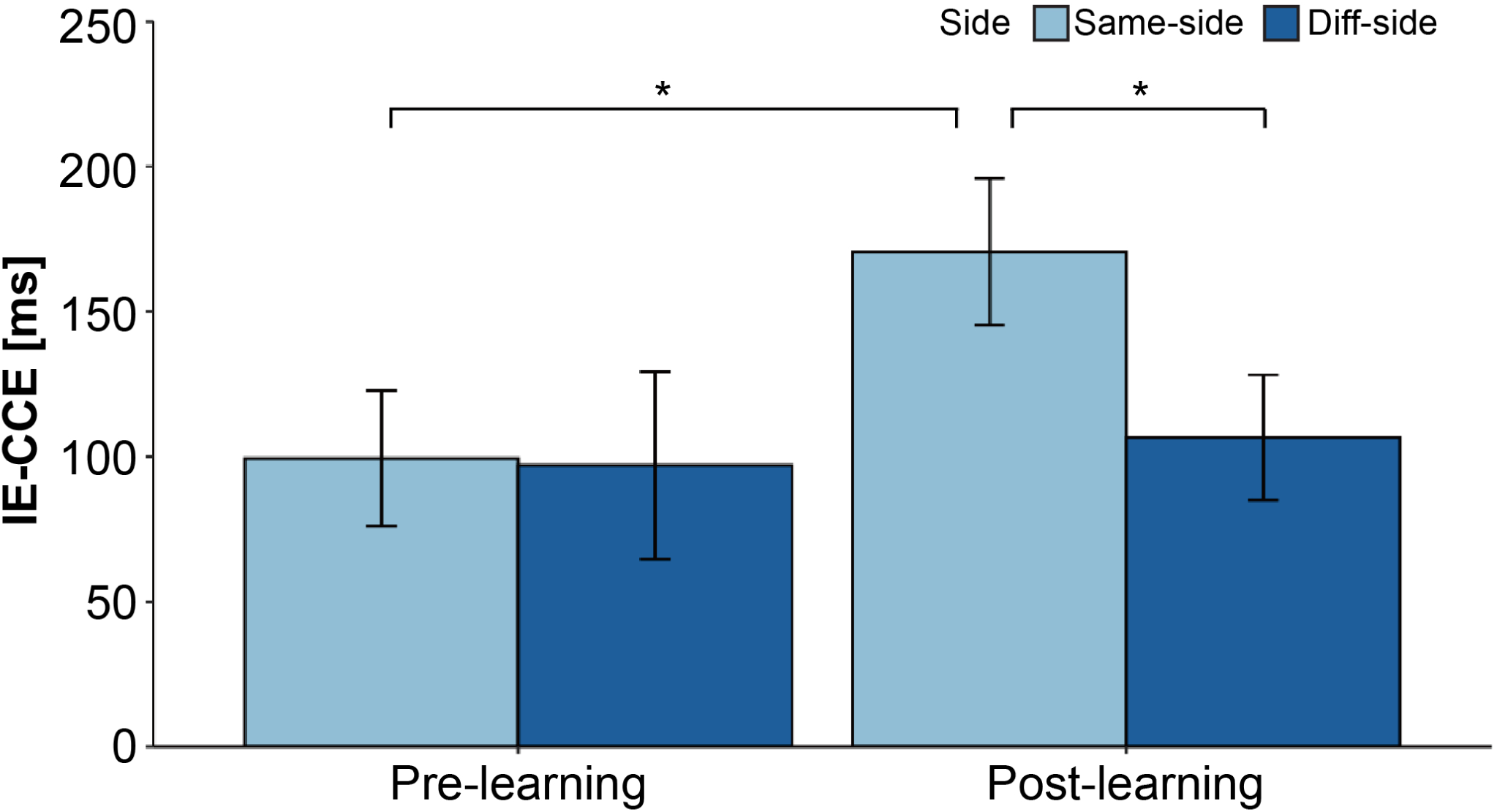
IE-CCE score. The IE-CCE is displayed for each *pre -*/*post-*learning, and the results are compared for the ipsilateral and contralateral conditions. Error bars represent the variability among the subjects for each condition. Pairwise comparisons by adjusting p-value for which a statistically significant difference (*ad j*.*p <* 0.05) can be confirmed are indicated by an asterisk (*). * … *ad j*.*p <* 0.05

### 7-Likert scale embodiment questionnaire

Questionnaires evaluating changes in embodiment toward an SRL (see Table 1) were conducted before and after the ball-touch learning task, and a statistical evaluation using the Wilcoxon-Mann-Whitney method was conducted by taking the averages among the subjects at each point in time. Figure 4 shows a box plot of embodiment questionnaire results per *pre-* and *post-*learning condition. The statistically significant differences were observed in Q1, “I felt as if the virtual robotic limbss/arms were my limbs/arms” [*p <* 0.001]; Q2, “It felt as if the virtual robot arms/limbs I saw were someone else’s” [*p <* 0.001]; Q3, “It seemed as if I might have more than two limbs/arms” [*p <* 0.001]; Q4, “It felt like I could control the virtual robot arms as if they were my own arms” [*p <* 0.001]; Q5, “The movements of the virtual robot arms were caused by my movements” [*p <* 0.001]; Q6, “I felt as if the movements of the virtual robot arm were influencing my own movements” [*p <* 0.001]; Q8, “I felt as if my arms were located where I saw the virtual robot arms” [*p <* 0.001]; and Q11, “At some point, it felt as if my real arms were starting to take on the posture or shape of the virtual robot arms that I saw” [*p* = 0.020 *<* 0.05]. As a trend, the participants tended to feel a sense of body ownership (Q1, 2, 3), a sense of agency (Q4, 5, 6), and a sense of self-location (Q8) after learning.

**Table 1.**
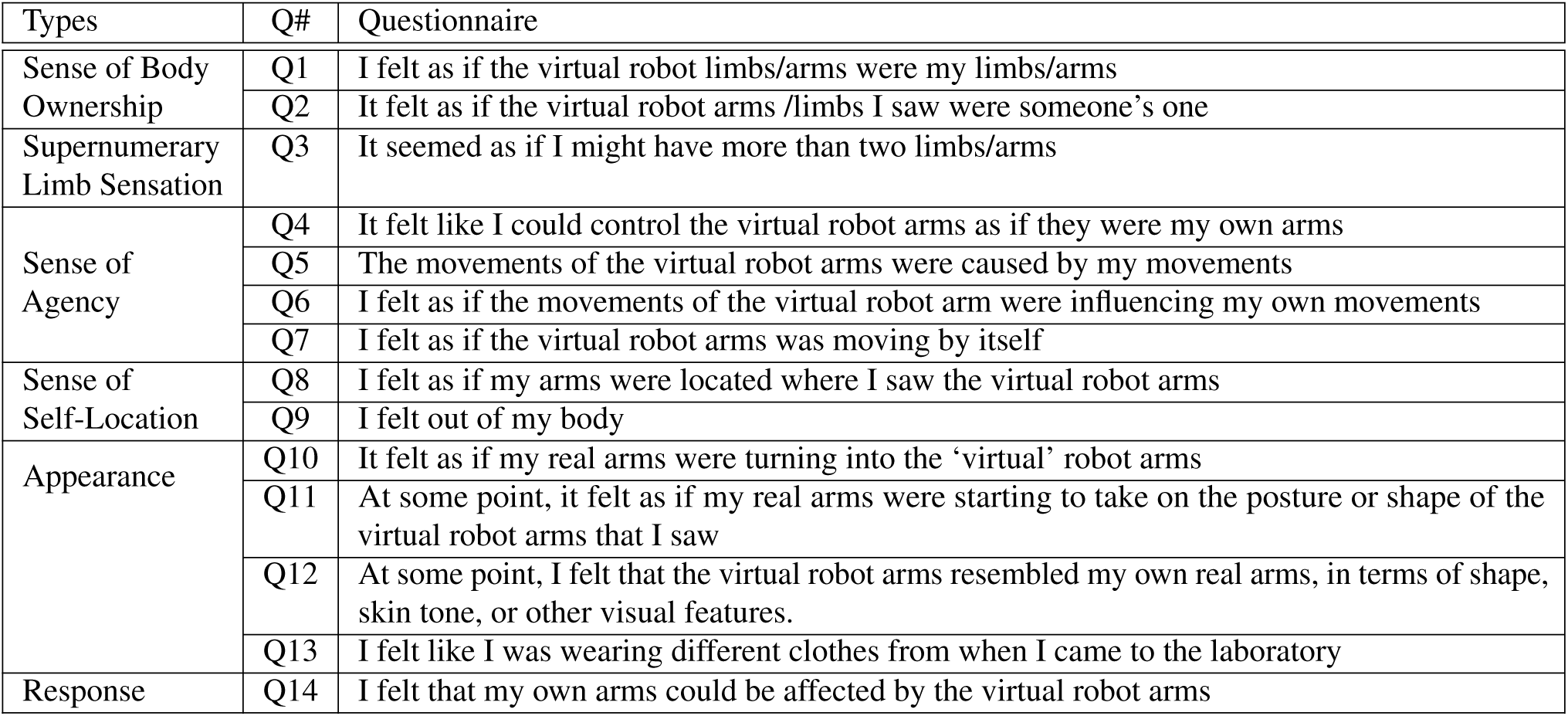
Embodiment Questionnaire. Subjects were asked to rate the questionnaire on a 7-Likert scale. Six types of questions were prepared: Sense of Body Ownership, Supernumerary Limbs Sensation, Sense of Agency, Sense of Self-location, Appearance, and Response.

| Types | Q# | Questionnaire |
| --- | --- | --- |
| Sense of Body Ownership | Q1 | I felt as if the virtual robot limbs/arms were my limbs/arms |
|  | Q2 | It felt as if the virtual robot arms /limbs I saw were someone’s one |
| Supernumerary Limb Sensation | Q3 | It seemed as if I might have more than two limbs/arms |
| Sense of Agency | Q4 | It felt like I could control the virtual robot arms as if they were my own arms |
|  | Q5 | The movements of the virtual robot arms were caused by my movements |
|  | Q6 | I felt as if the movements of the virtual robot arm were influencing my own movements |
|  | Q7 | I felt as if the virtual robot arms was moving by itself |
| Sense of Self-Location | Q8 | I felt as if my arms were located where I saw the virtual robot arms |
|  | Q9 | I felt out of my body |
| Appearance | Q10 | It felt as if my real arms were turning into the ‘virtual’ robot arms |
|  | Q11 | At some point, it felt as if my real arms were starting to take on the posture or shape of the virtual robot arms that I saw |
|  | Q12 | At some point, I felt that the virtual robot arms resembled my own real arms, in terms of shape, skin tone, or other visual features. |
|  | Q13 | I felt like I was wearing different clothes from when I came to the laboratory |
| Response | Q14 | I felt that my own arms could be affected by the virtual robot arms |

**Figure 4.**
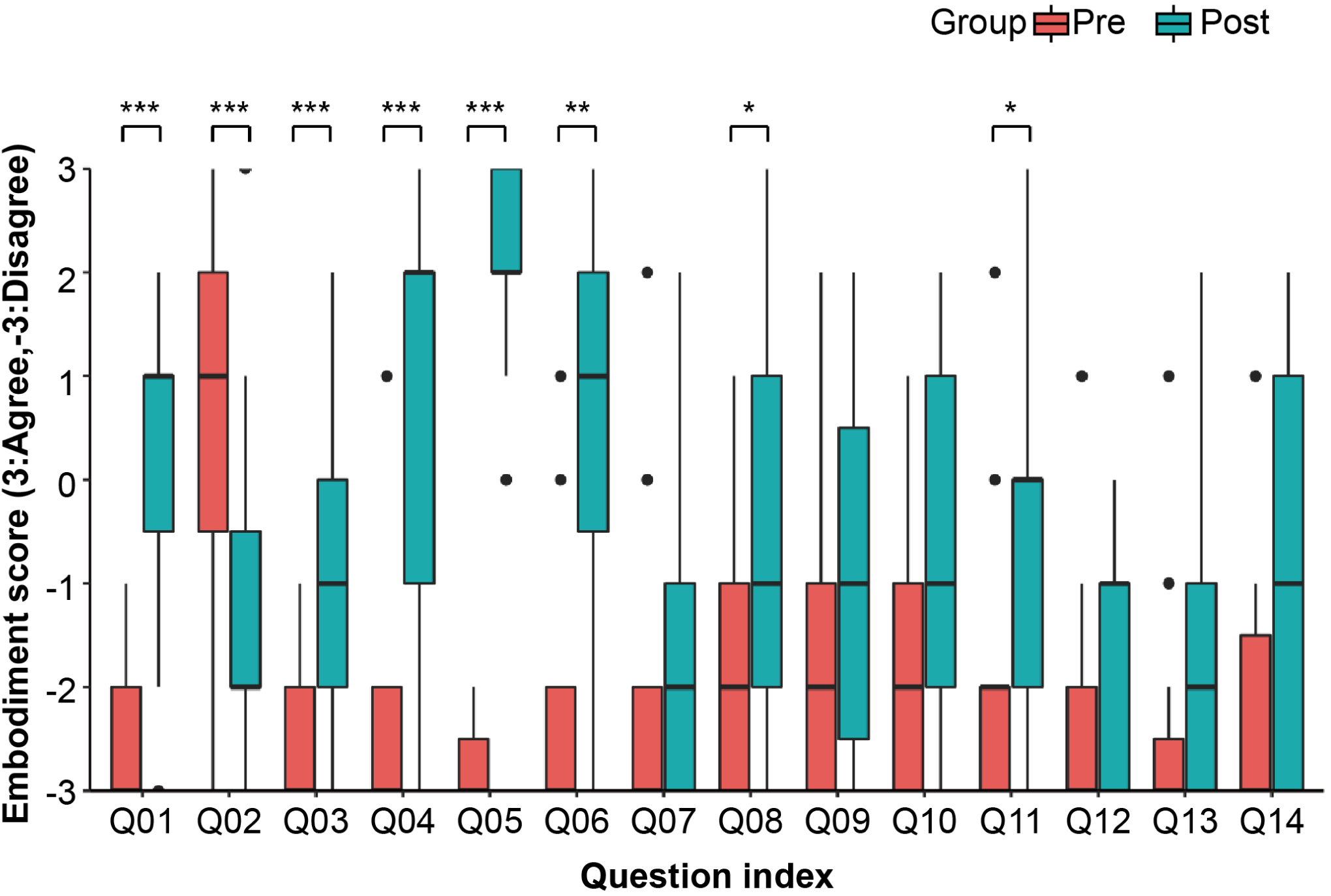
Embodiment Questionnaire Results. 7-Likert scales were used to collect responses to each question in each of the *pre-* and *post-*learning stages. Bar graphs are shown for each condition. The results are shown as boxplots for each condition. Asterisks (*) indicate statistically significant differences between the *pre-* and *post-*learning conditions. * … *p <* 0.05, ** … *p <* 0.01, *** … *p <* 0.001

### Correlation between embodiment questionnaire result and IE-CCE score

We focused on the relationship between IE-CCE scores and subjective evaluations of the embodiment of supernumerary limbs under each lateral condition before and after learning, where statistically significant differences were found. In this study, we focused on the amount of change and conducted a correlation analysis by taking the difference in each condition before and after learning. There were only 15 subjects in this study, and the sample size was insufficiently large to satisfy the normality required for a correlation analysis; thus, the analysis was conducted using a bootstrap method to theoretically satisfy the normality. As a result, there was a positive correlation between the IE-CCE score and the amount of change in the subjective evaluation Q3, “It seemed as if I might have more than two limbs/arms,” under the ipsilateral condition [adjusted *R*^2^ = 0.41, *F*(1, 1998) = 1380, *p <* 0.001] (see Figure 5). There was also a positive correlation between Q1, “I felt as if the virtual robot limbs/arms were my limbs/arms,” which refers to the sense of body ownership toward an SRL, and Q4, “It felt like I could control the virtual robot arms as if they were my own arms,” which refers to the sense of agency toward an SRL [adjusted *R*^2^ = 0.32, *F*(1, 1998) = 937, *p <* 0.001] (see Figure 6).

**Figure 5.**
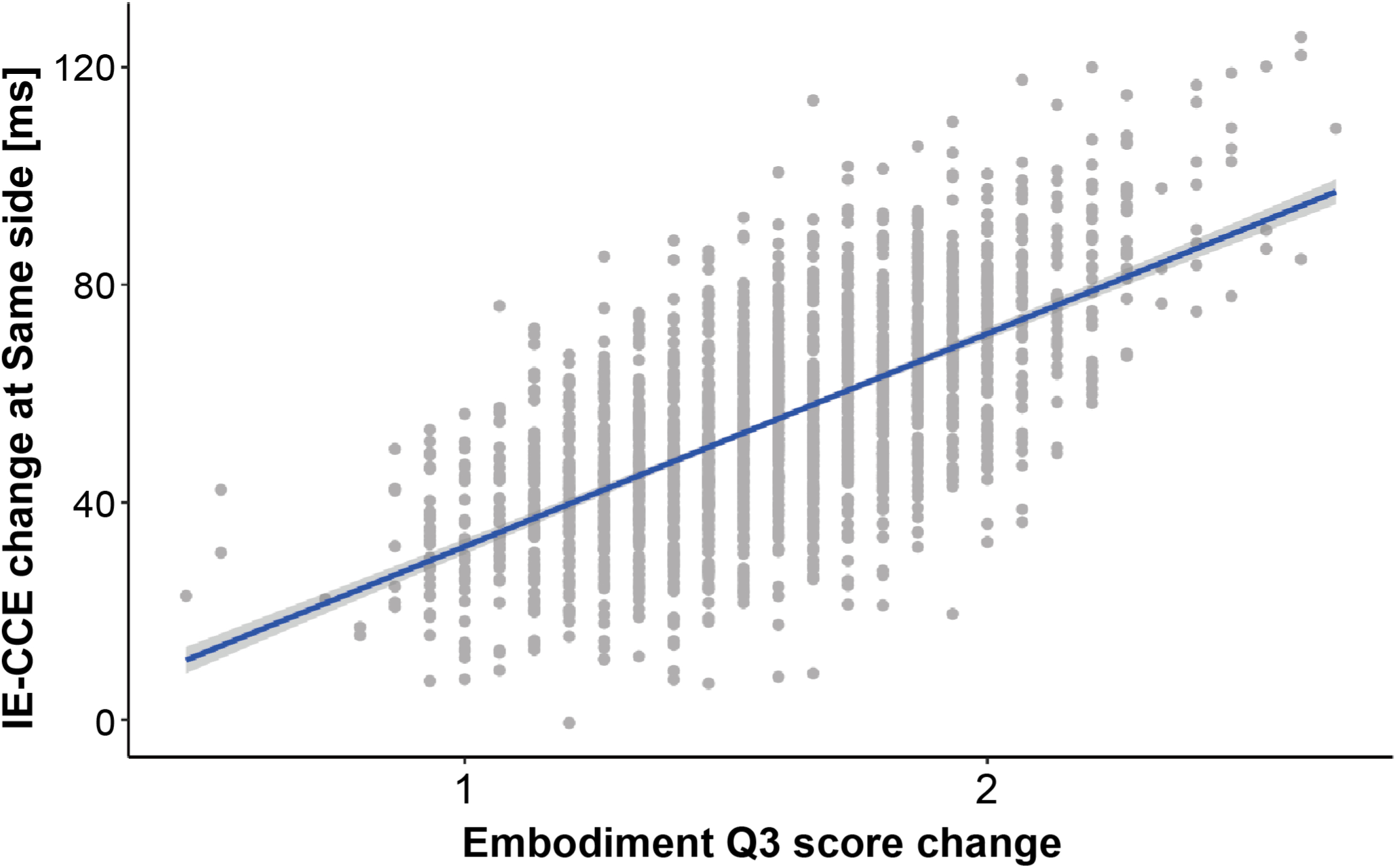
Correlation between the change in IE-CCE under the same-side condition and the change in response to the embodiment score Q3 before and after learning. Dots indicate points resampled by the bootstrap method, and linearly approximated lines and 95% confidence intervals are shown as the shadows of the regression lines.

**Figure 6.**
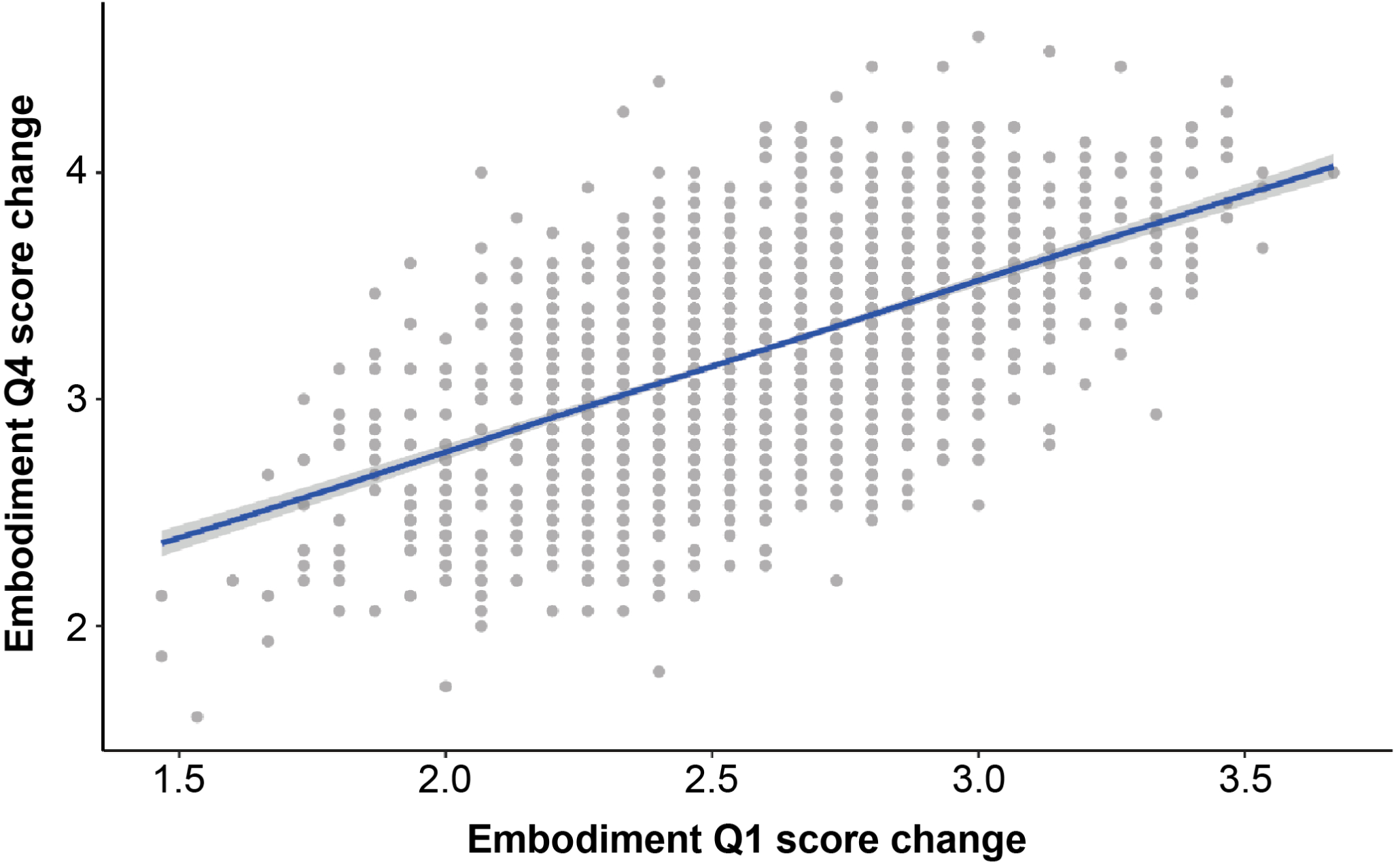
Correlation between the changes in response to the embodiment score Q1 and Q4 before and after learning. Dots indicate points resampled by the bootstrap method, and linearly approximated lines and 95% confidence intervals are shown as shadows of the regression lines.

## Discussion

In this study, we conducted experiments in a VR environment to clarify the embodiment of the SRL system and analyzed the following four aspects: (1) the duration time required for the learning task, (2) the IE-CCE score before and after learning the SRL manipulation, (3) the questionnaire score of the embodiment against the SRL, and (4) the correlation between (2) and (3). The results showed that the duration time required for each set of ball touch tasks became shorter for each set, and there was no significant difference in the time required between the last two sets. At the beginning of the task, it took time to learn the operation of the SRL system; however, in the latter half of the task, learning progressed and the time required was saturated in the third set. In the CCT, there were different trends in the IE-CCE scores of the same and different lateral sides before and after learning under the same lateral conditions, confirming cognitive changes owing to the visuotactile integration of the SRL. In addition, the 7-Likert scale questionnaire scores of embodiment changes toward the SRL showed that the sense of body ownership of the SRL, the sense of agency toward SRL manipulation, and the sense of self-location toward the SRL placed in the VR space significantly increased after learning the SRL manipulation. In addition, the correlation between IE-CCE score and the subjective evaluation of the embodiment to the SRL was examined. A positive correlation was found between the change of IE-CCE score under ipsilateral conditions and the amount of change in the supernumerary limb sensation. In addition, there was a positive correlation between the amount of change in the subjective evaluation of the SRL and the amount of change in the sense of body ownership and the sense of agency subjectivity toward the SRL.

### Imposing a sufficient number of tasks to become used to manipulating supernumerary robotic limbs in VR

The duration time required for the ball touch task per set was not statistically significantly different between the final two sets, suggesting that the subjects were able to become accustomed to manipulating the SRL through this task. In the interviews conducted after the subjects had learned to manipulate the SRL, the following comments were made: “The manipulation itself was fun,” “I was able to manipulate the SRL in the latter half of the second, third, and fourth sets without any particular awareness or cognitive load,” “I felt like my body naturally learned how to operate it,” “In the latter half, I was able to grasp the tempo and work on it like a rhythm game,” and “At first, I tried to manipulate the SRL with my hands, but I gradually got used to manipulating it with my feet.” These comments were supported by the fact that the total learning time, including breaks, was approximately 17–18 min. By contrast, there were some comments on the design of the SRL, such as “I felt confused because the movements of the SRL were contrary to the physical movements for manipulating them.” Others commented on the operation within the VR environment, saying, “It was the first time for me to experience VR, and I was a little tired.” Despite these comments, the duration time per set was saturated in the latter half of the final two sets.

### Embodiment assessment for supernumerary robotic limbs on VR-environment

The embodiment assessment of supernumerary limbs in a VR environment was conducted using the following three embodiment indices from a cognitive science perspective: (1) sense of body ownership of the SRL, (2) sense of agency in manipulating the SRL, and (3) sense of self-location in relation to the SRL placed in the VR space. In all of these cases, the changes in the subjective evaluations of embodiment before and after learning suggested that the SRL system was embodied. Although questions Q1, “I felt as if the virtual robot limbs/arms were my limbs/arms,” and Q2, “It felt as if the virtual robot arms/limbs I saw were someone else’s” are both evaluations of body ownership, they are contradictory questions. Both questions were inversely proportional and statistically significant, suggesting that the subjects felt a sense of body ownership toward the SRL. Q4, “It felt like I could control the virtual robot arms as if they were my own arms,” Q5, “The movements of the virtual robot arms were caused by my movements,” and Q6, “I felt as if the movements of the virtual robot arm were influencing my own movements,” were all statistically significant. By contrast, there was no statistically significant difference in Q7, “I felt as if the virtual robot arms were moving by themselves.” Therefore, it can be said that the subjects could feel a sense of agency toward SRL manipulation after learning. There was a statistically significant increase in the score of question Q8, “I felt as if my arms were located where I saw the virtual robot arms.” This suggests that the subjects could feel a sense of self-location toward the SRL placed in the VR space. A correlation analysis of the changes in the subjective evaluations of embodiment toward the SRL also showed a positive correlation between the sense of body ownership of the SRL and the sense of agency in the SRL manipulation. In light of these results, it can be considered that the subjects were able to embody the SRL after learning.

By contrast, there was no statistically significant difference after learning the SRL manipulation in Q10, “It felt as if my real arms were turning into ‘virtual’ robot arms,” and Q12, “At some point, I felt that the virtual robot arms resembled my own real arms, in terms of shape, skin tone, or other visual features.”

### Supernumerary limb sensation

In the present experiment, question Q3, “It seemed as if I might have more than two limbs/arms,” was handled differently from previous studies. In evaluations of rubber hand illusions and avatars in terms of sense of body ownership, this item has been regarded as a negative factor (i.e., no sense of body ownership)^51^. However, in our experiment paradigm, the interpretation is different because the SRLs are presented at the same time as the limbs of the avatar, which are mirrored to the movement of innate arms, and it is measured whether the sense of supernumerary limbs ownership is generated (supernumerary limb sensation). After learning the SRL manipulation, there was a statistically significant increase in the score, indicating that the subject may have learned the supernumerary limb sensation.

### Possibility that PPS occurred around supernumerary robotic limbs in VR

The results of the CCT showed that visuotactile integration toward the SRL caused perceptual changes, suggesting the possibility that a PPS occurred around the SRL. In the CCT, it is generally known that when a task-irrelevant visual distractor and tactile stimuli match (i.e., visuotactile stimuli appear at the same height), the response time is faster and the response accuracy is higher than when they do not match (i.e., visuotactile stimuli appear at different heights). This visuotactile interaction is a characteristic of the visual distractor stimuli when they are present within the vicinity of the body, and this characteristic is diminished when the visual distractor stimuli are presented far from the body. In the present study, we evaluated whether a strong integration of visual and tactile sensations, a typical feature of body perception, occurred within the vicinity of the body for the SRL, and found statistically significant increases in the IE-CCE scores before and after learning and in the IE-CCE scores on the same or different sides under the same lateral conditions. This result can be interpreted as a strong indication of post-learning visuotactile integration around the SRL, suggesting that PPS can occur. This is not only based on a statistical analysis, but also from interviews with the subjects conducted after the experiment, in which it was stated that “After learning, there were many cases of confusion when visual and tactile sensations were different in the CCT. In particular, I felt a stronger visual stimulus.”

### Embodiment of supernumerary robotic limbs in VR

In the present experiment, voluntary actions were encouraged to allow the subjects to learn to manipulate the supernumerary robotic arms, which is a different paradigm from the conventional rubber-hand illusion task based on passive actions and stimuli. In addition, our SRL is capable of arbitrary movements independent of an innate arm. Therefore, compared to the supernumerary hand illusion^39,40^, which is an extension of the rubber hand illusion used in previous studies, it is possible to evaluate the sense of agency toward the extra limb movement, and to conduct a more direct evaluation of the minimal self^20^. In the case of the ball touch task, the duration required to touch the ball in the second half of the task was significantly shorter than that in the first half. This may indicate that the active manipulation of the SRL facilitates the update of the body schema and stabilizes the motion plan of the SRL.

In this study, we examined the correlation between the IE-CCE score and subjective evaluation of the embodiment to the SRL. The correlation between the change in IE-CCE score under the ipsilateral condition and the subjective evaluation change of the embodiment to the SRL was positive. This suggests that there is a correlation between a supernumerary limb sensation and the generation of a PPS. In contrast to the “substitution” of embodiment to the extra limbs, which has been explained by the rubber hand illusion generated by passive stimuli, in the present study, the “addition” of embodiment was generated through learning and manipulation of the SRL with a voluntarily movable supernumerary limb. In other words, there is a possibility that the sense of supernumerary limb ownership could be explained by the simultaneous generation of the supernumerary limb sensation and PPS. In addition, bodily self-consciousness (BSC)^52^ is a cognitive representation of one’s own body, and perceptions of embodiment and PPS have been cited as necessary conditions for achieving BSC. The fact that these correlations were observed in our study suggests that the SRL system was incorporated as a self-body in the context of the BSC.

### Cognitive transparency for supernumerary robotic limbs system

It is important to suppress the cognitive load (i.e., cognitive transparency) during the operation of the SRL system working as a new body part. The human–robot coupling system^53,54^ has been proposed as a control strategy for the entire supernumerary limb robotic system. In this system, there are components that recognize the operator’s intentions and mechanisms to return the expected feedback. However, there is no mention of how the cognitive science elements of the man–machine interface should be present in the system. When there is a concern that when the cognitive workload is high, it may interfere with the accuracy of the operation^8^; thus, it is desirable to suppress the cognitive load. Therefore, it is also desirable that the man–machine interface should be designed with a cognitive load.

The indices and evaluation paradigms verified in this study may be utilized in the design and evaluation of SRL systems in the future. Particularly in terms of perceptual and cognitive changes, we examined the possibility of PPS using the CCE score and assessed the subjective evaluation of the embodiment of the SRL system in a VR space. This can be considered an evaluation of an example of many SRL systems. This evaluation approach can be used to compare the ease of embodiment of other SRL systems when examining and evaluating their potential for embodiment.

While gender effects to cause the biased result in VR studies was reported^55^, 90% of the participants in this study were male. Thus, well-balanced gender ratio should be required for experimental design in future study.

### Required evaluation of cooperative work using innate and supernumerary robotics limbs

In this study, we did not evaluate the task using the cooperative work of innate limbs and SRL; therefore, an additional validation is needed to further discuss the transformation of body schema. In a previous study, it was claimed that rubber hands can be incorporated into a body image and body schema in the rubber hand illusion system; however, only one of the limbs (rubber or innate hand) can be incorporated into the body schema, and an exclusive relationship has been reported, particularly in the body schema^40^. By contrast, in the robot system developed in this study, while the innate limbs are presented as avatars, the robotic arm itself can perform arbitrary movements independent of the innate arms, suggesting the possibility of SRL embodiment in terms of the three elements of body ownership, sense of agency, and sense of self-location. Considering this, our SRL system is clearly different from the experiment and evaluation systems for rubber hand illusions. However, because we have not evaluated the system in cooperation with the innate limb, we cannot say whether the body schema will be exclusively transformed or whether a body schema incorporating both the innate limb and the supernumerary robotic limb will be constructed.

### Required evaluation of perception change of innate feet mapped using supernumerary robotic limbs

To discuss in depth the generation of a supernumerary limb sensation, it is necessary to evaluate the perceptual change of the feet mapped with the SRL. In this study, we assume that the sense of body ownership and the sense of agency against the innate limbs are maintained because the SRL is presented at the same time as the avatar mirroring the innate limbs. Based on this, the subjective evaluation of embodiment question Q3, “It seemed as if I might have more than two limbs/arms,” was interpreted as a supernumerary limb sensation because its treatment was different from the experimental paradigm of previous studies on the rubber hand illusion and avatar embodiment. To refer to this more rigorously, we can describe the relationship between the supernumerary limb sensation and perception in the mapped innate limb by evaluating the perceptual changes in the innate feet.

### Future works and conclusion

The findings of this study are not limited to an evaluation of the embodiment of the SRL, and also suggest the possibility of further subdividing the discussion on the embodiment of tools used in cognitive science. Traditionally, the “substitution” and “stretching” of perceptual changes when using tools such as knives, scissors, and canes have been highlighted. However, we believe that with the advent of SRL systems that enable the addition of body parts and functional additions, we will be able to capture perceptual changes that are an “addition.” The possibility of the supernumerary limb sensation reported in this paper is an example of capturing “additional” perceptual changes. In addition to “substitution” and “stretching,” an “addition” can be included in the study of perceptual changes in the embodiment of tools, which may lead to further subdivision in cognitive science discussions. Consequently, it will be possible to discuss this in research fields such as neuroscience.

As a future prospect, we believe that it will be necessary to capture the transformation of a perception using a neuroscience approach, in addition to the aforementioned cooperative behavior experiments between the innate limb and the extra-limb robotic system, and to verify the perceptual changes in the innate limb mapped to the SRL system. Although previous studies have evaluated neural representations for extra fingers^56^, it is unclear how neural representations differ depending on the area added as a supernumerary limb. In addition, there are many items that need to be considered, such as the learning task and the time required to wear the SRL system. In addition, from the standpoint of cognitive neuroscience, it will be important to investigate the mechanisms and dynamics of the supernumerary limb sensation and extra-limb possession sensation reported herein to explore the limits of human plasticity and design SRL systems.

In this paper, we suggest that the participants might feel that they have acquired a new body part that is different from their own body part through the use of an SRL presented in a VR space. There was a correlation between the PPS generation around the SRL and the sensation of gaining a new body part (supernumerary limb sensation), which was used to explain the possibility of embodiment of the SRL system. In addition, we suggest the possibility of subdividing perceptual changes in the embodiment of tools to facilitate cognitive science discussions.

## Methods

### Participant recruitment, experiment logistics, and experiment procedure

Sixteen healthy subjects (including two females, ranging in age from 21 to 27, with an average age of 22.9) participated in this experiment as paid volunteers, regardless of gender, handedness, or footedness. One participant had to stop the experiment because of equipment trouble, and therefore was not included in the analysis. We used data from 15 participants in the analysis. Referring to previous studies, Aspell et al. CCT in Study 1 [*F*(1, 12) = 11.3]^57^. The sample size calculated based on the significance level [*α* = 0.05] was 14 participants, which met the required sample size^58^. All participants used in the experiment were confirmed to have normal or corrected-to-normal stereo vision. The experiment was conducted after approval by the Ethics Committee of the Research Center for Advanced Science and Technology, the University of Tokyo, and written informed consent was obtained from all participants prior to participation. All experiment procedures described below were approved by the same ethics committee, and were conducted in accord with the guidelines of the Declaration of Helsinki^59^ and the approved procedure by same ethics committee.

### Experimental setup for VR-based supernumerary robotic limbs system

The participants in the experiment saw that they were placed in a virtual space through a head-mounted display (HMD, VALVE INDEX, presenting stereoscopic images with a resolution of 2880×1600). The virtual space was constructed using Unity3D (FPS fixed at 60) and was run on a Windows PC (Razer BLADE15, 2.6-GHz Intel Core i7-9750H, 16 GB of RAM, and an NVIDIA RTX2080). The participants were positioned in a VR environment so that they could see the avatar’s body including the limbs and the supernumerary robotic arms from a first-person perspective.

The SRL system operated in a VR environment was designed based on Sasaki et al.’s system design using limb mapping^3^. The supernumerary robotic arms consists of seven degrees of freedom: the shoulder joint, upper arm joint, elbow joint, lower arm joint, wrist joint, and end-effector. The avatar moves in conjunction with six points in total, i.e., the tracking points on the HMD, the VALVE INDEX controller, and the VIVE trackers attached to the waist and toes. In particular, the movement of the VIVE tracker at the feet is linked not only to the avatar’s feet, but also to the robotic arms. The VIVE tracker information, together with the distance to the end-effector, is set as the target position for inverse kinematics in the vertical and horizontal movement and rotation of the robot arm, and thus the robot arm can move within the range required for the adaptation task. This enables the robotic arms to move as desired. Depending on the manipulation position, it may be necessary to move the linked leg in the opposite direction and extend the hand part of the robotic arm forward from the avatar’s perspective.

To provide tactile feedback when the robotic arm touches the ball presented in a virtual space, we used Arduino to link Unity3D with an oscillator. The transducer is a small vibro-transducer (Acouve Lab’s Vp210), and the vibration frequency was set at 200 Hz, which is a frequency band with high sensitivity to human vibration detection^60^. The back of the subject’s foot vibrates against the back of the hand of the robotic arm, and the sole of the subject’s foot vibrates against the palm of the robotic arm. Four transducers, including the left and right, were attached to the subject’s feet. During the task, white noise was continuously played to block out sounds from the outside world, and the auditory stimuli were controlled to be constant.

To collect responses to the stimuli as quickly and accurately as possible during the CCT, we used a vertical ergonomically designed mouse (Delux M618-PLUS, polling rate = 500 Hz, USB wired connection).

### Experiment procedure

First, calibration was conducted to place the subject in the virtual space as intended. The subject was then equipped with the equipment mentioned in the experiment setup. As soon as they were ready, the subjects worked on the CCT without practicing or learning to use the supernumerary robotic arms. Immediately afterwards, they answered a questionnaire related to their subjective evaluation of embodiment. To answer the questionnaire, the HMD was removed once. After answering the questionnaire, the participants engaged in a ball-touch task as an adaptation task. Immediately after the series of adaptation tasks were completed, the participants tackled the crossmodal matching task again and answered the questionnaire. Finally, they were interviewed to collect their impressions regarding the entire experiment. The details of each item are described below. Although we did not set a time limit, from the preparation to the end of the experiment took approximately 90 min for all subjects.

### 7-Likert embodiment questionnaire

The subjects were asked to complete an embodiment questionnaire administered immediately after the CCT. The subjects were then asked to rate the questionnaire on a 7 Likert scale ranging from 3 (strongly disagree) to +3 (strongly agree). Six types of questions were prepared: body ownership, supernumerary limb sensation, sense of agency, sense of self-location, appearance, and response (see Table 1)^51^. In this experiment system, unlike conventional experimental paradigms such as the rubber hand illusion, we are trying to add new body parts rather than substitute them; therefore, the interpretation of Q3, “It seemed as if I might have more than two limbs/arms,” is different. In this study, we collected subjective evaluations of the supernumerary limb sensation for extra body parts that can be arbitrarily moved. Some questions were prepared to ask if the distinction was made between participant’s arm and robot arm.

### Crossmodal Congruency Task

In the CCT, the participants make discriminative judgments regarding the presentation position (up or down) of tactile stimuli while ignoring task-irrelevant visual distractor. During the CCT, the subjects were instructed to look at the gazing point through the HMD throughout the task and were asked to respond immediately after the tactile stimulus was presented. The subject had to answer with the upper button of the mouse when the vibrator attached to the toe presented the stimulus to the top of the foot, and with the lower button when the stimulus was presented to the sole of the foot. The latency between the visual and tactile stimuli (stimulus onset asynchrony, SOA) was designed to be 33.3 ms, and the system latency was confirmed to be 16.7 ms at maximum from the logs recorded during the experiment. The maximum system delay was confirmed to be 16.7 ms from the logs recorded during the experiment (i.e., the actual SOA was 33–55 ms). It is known that when the SOA is less than 100 ms, multisensory integration (MSI) contributes to the CCE score, and when the SOA is outside this range, the CCE score is mainly influenced by extrinsic attention and priming effects^61–63^. The tactile stimuli were presented 1 s after the cue was presented, and the time between the tactile stimulus and the mouse’s response was recorded as the RT. The next cue was presented 3 s after the subject’s response, and the sequence was set up such that the next cue was presented again.

The combination of visual and tactile stimuli was determined such that the visual stimuli satisfied the conditions of (*ipsilateral*/*opposite*) and (*congruent*/*incongruent*) for the tactile stimulated foot (*right*/*left*). There were eight patterns in total. The order of the stimulus presentation was randomized, and six sets of combinations were presented per session, with a break after each session and an immediate transition to the next session if not required. The subjects conducted 192 trials (8 *patterns* × 6 *sets/session* × 4 *sessions*) before and after manipulating the SRL.

The CCE was calculated from the difference in the RT between *incongruent* and *congruent* trials. To account for the fact that some responses may be incorrect, the IE, which is the percentage of correct responses divided by the average reaction time under a particular condition, was also used as an index when calculating the CCE^34,49^.

### Ball-touch task as learning of supernumerary robotic limbs manipulation

We set up a session to learn how to use the supernumerary robotic arms without a time limit. The subjects were asked to touch the upper ball with the back of the hand of the robot arm and the lower ball with the palm of the robot arm. When the subject touched the ball, it disappeared, and tactile stimuli were simultaneously presented to the corresponding foot. The ball was then immediately presented to a different location, and the subject conducted this series of tasks a total of 400 times. A break was provided every 100 trials to allow for subject fatigue. The order of the appearance of the balls was randomly determined. The time until the next ball touched was measured, and the positions of the trackers on the left and right toes were also recorded. In contrast to the CCT, there was no restriction on the point of view, and a display counting the number of times the ball was touched was shown at a different location from the ball.

## Data availability

The datasets generated and analyzed during the current study are available from the corresponding author on reasonable request.

## Acknowledgements

This research was supported by JST ERATO Grant Number JPMJER1701 (Inami JIZAI Body Project).

## Author contributions statement

K.A., H.S., M.F., S.U., and M.K. conceived and designed the experiments. M.F. developed the base system. K.A. collected and analyzed the data. H.S., S.U., and M.K. contributed to the preparation of the manuscript. K.A. obtained and created all images, drawings, and photographs. K.A. and H.S. created a movie attached as supplemental material. All authors reviewed the manuscript.

